# Tuneable hydrogel stiffness in a 3D *in vitro* model induces epithelial to mesenchymal transition in MCF7 but not MDA-MB-231 breast cancer cells

**DOI:** 10.1101/2023.06.29.546799

**Authors:** JA Wise, MJ Currie, TBF Woodfield, KS Lim, E Phillips

**Affiliations:** Mackenzie Cancer Research Group, Department of Pathology & Biomedical Science, University of Otago Christchurch, Aotearoa New Zealand; Christchurch Regenerative Medicine and Tissue Engineering (CReaTE), Department of Orthopaedic Surgery and Musculoskeletal Medicine, University of Otago Christchurch, Aotearoa New Zealand; School of Medical Sciences, University of Sydney, Sydney, Australia

**Keywords:** GelMA hydrogel, breast cancer, 3D model, epithelial to mesenchymal transition (EMT)

## Abstract

The study of *in vitro* models of breast cancer is crucial for understanding and treating the malignancy in patients, with 3D *in vitro* models providing researchers with more biomimetic systems to overcome limitations of current to 2D cultures and *in vivo* animal models. *Ex vivo* patient tissues have shown that malignant breast tissues are stiffer than healthy or benign tissues, and that the stiffness corresponds with increasing tumour grade. Stiffening of the breast tumour environment alters tumour cell phenotype and facilitates tumour progression, invasion and metastasis. Better understanding of the relationship between extracellular matrix stiffness and breast cancer cell phenotype, and how that is important in the initiation of metastasis, should lead to designing 3D models that mimic the breast tumour microenvironment at different stages of breast cancer progression.

This study investigated phenotypic response of two breast cancer cell lines that are representative of clinical breast cancer subtypes (MCF7, Luminal A; MDA-MB-231, Triple Negative Breast Cancer) in gelatin-methacryloyl (GelMA) hydrogels of varying stiffness. A visible light photoinitiation system was adopted to provide a tuneable photocrosslinking platform to systematically control hydrogel stiffness and tumour microenvironment. This allowed rapid fabrication of biocompatible hydrogels supporting high cell viability over long-term culture.

The impact of a clinically relevant range of microenvironmental stiffness on breast cancer cell behaviour and phenotype was examined over a 21-day culture period using GelMA hydrogels. Results showed that MCF7 cells cultured for 21 days in high stiffness hydrogels (10 wt%; 28 kPa) responded by downregulating the epithelial marker E-cadherin and upregulating mesenchymal markers N-cadherin and Vimentin, whereas MDA-MB-231 cells showed no changes in EMT-markers when cultured in hydrogels of corresponding stiffness (10 wt%; 33 kPa). Culturing both cell lines in soft hydrogels (5 wt%; 11 kPa) maintained their phenotype over 21 days, highlighting the importance of controlling hydrogel mechanical properties when studying breast cancer cell phenotype.

## Introduction

The tumour microenvironment (TME) plays a significant role in breast cancer progression^1–3^. Breast cancer cells and the TME interact with each other, resulting in a supportive “reactive tumour stroma”^3–5^. The TME is comprised of various cell types as well as acellular components, such as the extracellular matrix and oxygen/nutrient gradients. The properties of the extracellular matrix have been shown to impact various aspects of tumour biology such as breast cancer cell adhesion, metabolism, proliferation, differentiation, migration, and therapeutic resistance^6–11^. The stiffness of the extracellular matrix is of particular significance, as it has been observed that human breast tumours are stiffer than healthy breast tissue, and increasing stiffness has been linked to higher tumour grade^12–16^. Mechanical testing of *ex vivo* patient tissues have shown normal fibro-glandular breast tissue has a stiffness of roughly 3.2 kPa, while invasive ductal carcinomas (IDCs) increase in stiffness with the grade of cancer (ranging from 10.4 to 42.5 kPa from low to high grade IDC, respectively)^12, 13, 17, 18^.

The development of 3D *in vitro* culture models for breast cancer offers a more authentic representation of the *in vivo* microenvironment than traditional 2D *in vitro* culture techniques^19–22^. With 3D *in vitro* models, cells are cultured in a microenvironmental matrix that closely resembles the *in vivo* tissue and directly impacts mechanical properties and cell-to-cell interactions^23–26^. These models are also more cost-effective and efficient for screening new therapeutic treatments than animal models^19, 27–29^.

GelMA hydrogels are an appealing biomaterial for 3D models due to their tuneable physico-mechanical properties^22, 30, 31^. For example, hydrogel stiffness can be altered by adjusting the amount of macromer added to the hydrogel, while maintaining the hydrogel’s ability to allow for the diffusion of liquids^22^. Research has shown that cells cultured in GelMA hydrogels exhibit high viability, metabolism, adhesion, and proliferation, as well as the formation of spheroids with some cell types^21, 22, 32–34^. However, there is limited research on the effects of GelMA hydrogel culture on breast cancer cells, with only a few studies conducted over a period of 14 days, and focusing mainly on cell viability, metabolic activity, and migration^30, 35–37^.

This study is the first to report the use of GelMA hydrogels crosslinked using a visible light photoinitiator system in conjunction with breast cancer cells. Herein, we describe using a photoinitiator system comprising Ru/SPS and visible light (400-450 nm) to rapidly fabricate biocompatible 3D hydrogels that allow better light penetration and maintain high cell viability^33, 38–43^. The visible light photo-crosslinking used in this study offers an alternative to commonly used UV crosslinking approaches, which have been shown to lead to chromosomal damage and reduced cell viability^33^.

The aim of this study was to investigate the effects of extracellular matrix stiffness on breast cancer cell phenotype and behaviour, using 3D visible light crosslinked GelMA hydrogels encapsulated with two different types of breast cancer cells: MCF7 cells, representative of Luminal A hormone responsive breast cancer subtype; and MDA-MB-231 cells, which are representative of the triple receptor negative breast cancer subtype. By comparing the response of these two breast cancer cell lines to clinically relevant hydrogel stiffness, this study offers insights into how breast cancer cells behave in response to extracellular matrix stiffness within the breast tumour microenvironment.

## Materials and Methods

### Cell culture

The human breast cancer cell lines MDA-MB-231 (HTB-26™) and MCF7 (HTB-22™) were obtained from the American Tissue Culture Collection (ATCC^®^; Manassas, VA, USA). The cells were cultured in high glucose Dulbecco’s Modified Eagle Medium (Gibco, USA) supplemented with 10% fetal bovine serum (Gibco, USA), 1% penicillin-streptomycin (ThermoFisher Scientific, USA) and 0.1% amphotericin B solution (Sigma-Aldrich, USA) under aseptic conditions in a humidified 37°C incubator with 5% CO₂. Experiments were performed within the first 28 passages of cell culture, and cell lines tested negative for mycoplasma contamination.

### GelMA hydrogel preparation

GelMA synthesis was carried out as previously described^44^. Briefly, a 10 w/v% solution was prepared by dissolving gelatin (from porcine skin) in 50°C 1x PBS solution and 0.6 g of methacrylic anhydride (MA) added per gram of gelatin. Following dialysis against deionised water, the solution was then pH adjusted to 7.4 and sterile filtered (0.2 μm pore size) before being frozen and lyophilised. This resulted in GelMA with a 58% DoF. To produce breast cancer cell-laden GelMA hydrogels, the lyophilized GelMA was dissolved in 1x PBS to produce 5%, 7.5%, and 10% wt% solutions and pelleted MDA-MB-231 or MCF7 breast cancer cells incorporated at the indicated final concentration of cells/mL. The photoinitiators Tris (2,2’-dipyridyl)dichloro-ruthenium(II) hexahydrate (Ru) and sodium persulfate (SPS) were added at 0.5 and 5mM, respectively, and the resulting solution pipetted into silicone moulds (1mm deep and 5mm diameter for biological studies, 2mm deep and 5mm diameter for mechanical testing). Crosslinking was performed using visible light (400-450 nm) with an intensity of 30 mW/cm² for 180 seconds (OmniCure S1500, Excelitas Technologies with a Rosco IR/UV filter). Cell-laden hydrogels were transferred to individual wells of a 48-well plate with 500 µL of cell culture media, which was refreshed every 3-4 days (Figure 1).

**Figure 1:**
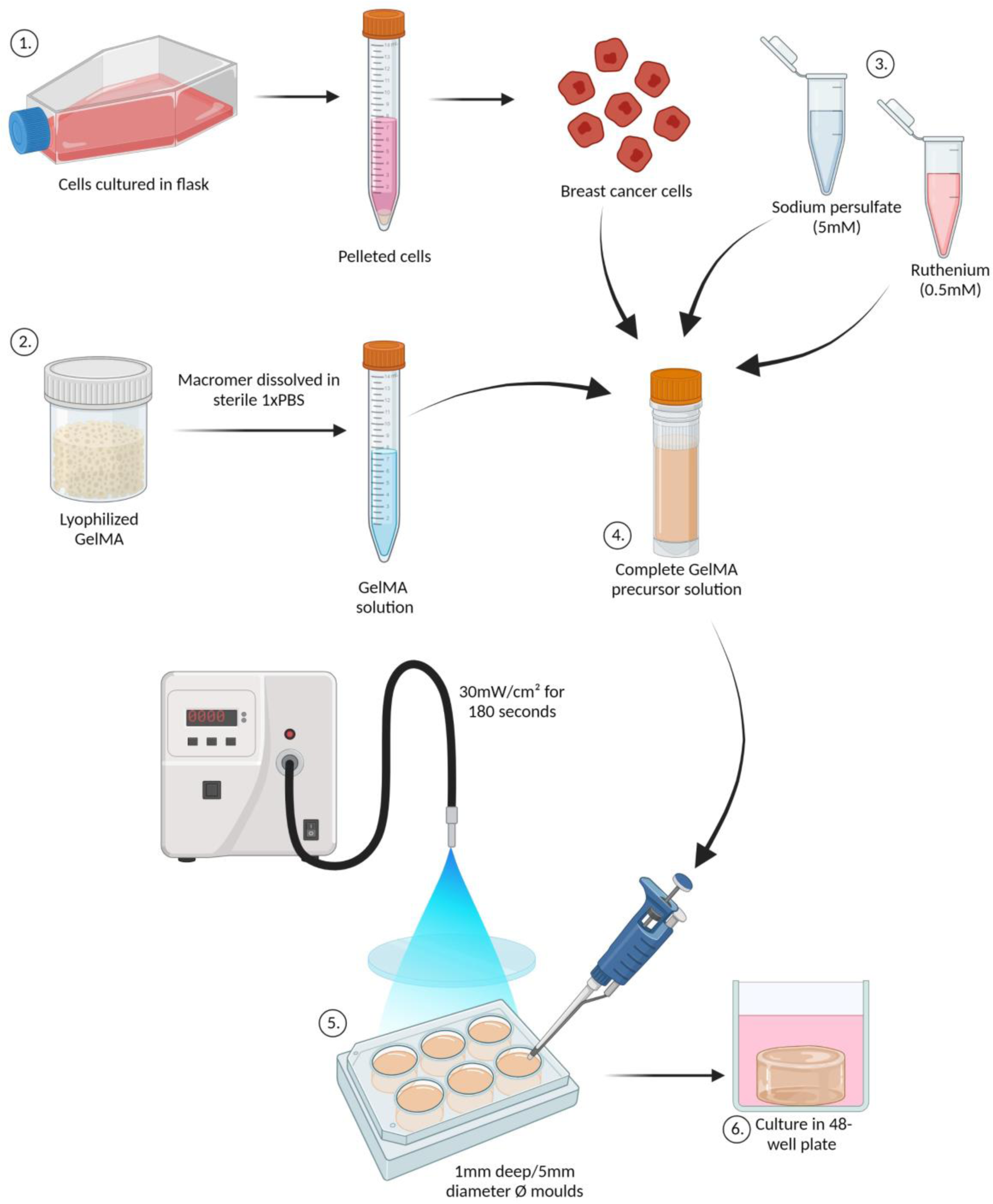
Schematic of GelMA hydrogel preparation with cell encapsulation. (1) Breast cancer cells were detached from the cell culture flask then pelleted to obtain the desired final concentration of live cells/mL, (2) Lyophilized GelMA macromer was weighed and dissolved in sterile 1x PBS solution to produce the GelMA solution of designed wt%, (3) Sodium persulfate (SPS) and ruthenium (Ru) were dissolved in sterile 1x PBS to produce solutions with final concentrations of 5 and 0.5 mM, respectively, (4) To produce the complete GelMA precursor solution; the dissolved GelMA solution, pelleted cells and photoinitiators (SPS and Ru) were combined and pipetted into 1 mm deep, 5 mm Ø moulds. (5) These were then crosslinked with 30 mW/cm² visible light for 180 seconds, before being transferred into individual wells of a 48-well plates with 500 µL cell appropriate media (6). Image created with Biorender.

### Analysis of GelMA hydrogel properties

Hydrogels were weighed immediately following crosslinking to obtain the initial mass (m initial, t0), then either incubated overnight at 37°C in a humidified cell culture incubator in 1x PBS, or immediately frozen in a −80°C freezer before freeze drying (m dry, t0). The hydrogels which were swollen overnight were then weighed again (m swollen), before being frozen in a −80°C freezer. All hydrogels were then freeze-dried and the final dry weight (m dry) measured. These values were used to calculate actual macromer fraction (1), initial dry mass (2), soluble fraction (%) (3) and mass swelling ratio (*q*) of hydrogels (4).

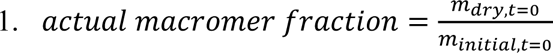

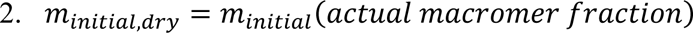

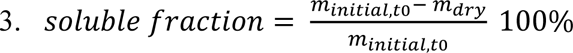

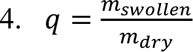

Hydrogels, 2 mm in height and 5 mm diameter, underwent mechanical compression testing using an MTS Criterion machine (Model C42) fitted with a 5 N load cell to determine compressive modulus. The compressive modulus (E) was calculated (equation 5) to produce a stress (σe) – strain (εe) curve for each hydrogel using the compression testing data, hydrogel height and cross-sectional area measurements. The slope of the linear region of the stress-strain curve determined the mechanical stiffness (compressive modulus) of each hydrogel. The 10-20% strain region of the curve was found to show the highest R2 values representing the linear region of these hydrogels and was used for all experiments.

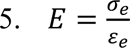

A Presens^®^ oxygen microsensor probe was used to measure oxygen levels in a step-wise manner throughout the depth of cell-free GelMA hydrogels. The oxygen probe microsensor was mounted in the head of a computer controlled 3D printer (Bioscaffolder, SYSEng) to allow for precise control over the z position of the probe tip within hydrogels. The sensor was positioned into the centre-bottom of the hydrogel and allowed to equilibrate for 1 minute, then retracted at 100 µm every 10 seconds (with 10 measurements per second) to provide step-wise measurements throughout the depth of the 1 mm hydrogel. Measurements are denoted as 0 µm at the top of the hydrogel and 1000 µm at the bottom of the hydrogel.

### Measurement of breast cancer cell viability, metabolic activity and DNA content

The study evaluated cellular response to 3D GelMA hydrogel culture by conducting Live/Dead staining to determine cell viability, AlamarBlue^®^ to measure metabolic activity, and CyQUANT^®^ to assess cellular proliferation on day 1, 7, 14, and 21 of the culture period.

Briefly, Live/Dead staining was performed with Calcein acetoxymethyl (Calcein-AM) and propidium iodide (PI). Hydrogels were washed with 1x PBS for 5 minutes, then incubated in Calcein-AM and PI (1 µg/mL) for 10 minutes. Hydrogels were then washed with PBS before fluorescent imaging on the Zeiss Axio Imager.Z1 Microscope while hydrated. ImageJ software was used to count the number of live and dead cells and calculate the percentage viability (%) (equation 6).

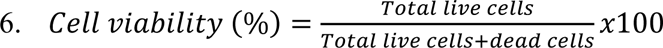

The AlamarBlue™ cell viability assay was used to assess the metabolic activity of encapsulated cells. Breast cancer cell laden hydrogels were incubated with 10% v/v Alamar blue reagent in cell culture media for 6 hours, after which the fluorescence intensity measured at excitation of 545 nm and emission at 590 nm with the VarioSkan™ plate reader. Results were normalised to negative controls and expressed as relative fluorescence units (rfu).

Breast cancer cell proliferation was measured using the CyQUANT™ direct cell proliferation assay. Cell-laden hydrogels were collected at indicated timepoints and washed with 1x PBS to eliminate residual phenol red from media. The hydrogels were then frozen at −80°C and subjected to freeze-drying. The lyophilized hydrogels were then digested in proteinase K solution (1 mg/mL in Tris-EDTA) at 56°C for 12 hours. The CyQUANT™ assay kit was used to quantify the DNA content of each hydrogel, following the manufacturer’s protocols. The experimental measurements were normalized to negative controls, and results were extrapolated from the DNA standard curve.

### Immunofluorescent staining

To prepare for staining, hydrogels containing cells were washed with 1x PBS and then fixed in 10% neutral buffered formalin at room temperature for 1 hour. The hydrogels were then frozen at −80°C in O.C.T. before being cryosectioned into 30 µm sections. The cryosections were simultaneously blocked and permeabilized in 3% w/v BSA and 0.1% v/v Triton™ X-100 before incubation with primary antibodies (Anti-Ki67; 1: 500, Abcam, USA) (Anti-E-cad/N-cad/Vimentin; 1:200, Cell Signalling, USA) (Rhodamine phalloidin dye, 6.6 µM, ThermoFisher Scientific, USA) at room temperature for 1 hour. The sections were then washed with 1x PBS-T before incubation with the secondary antibody (Goat Anti-Rabbit IgG (Alexa Fluor® 488); 1:1000, Abcam, USA) overnight at 4°C. The sections were counterstained with 4′,6-diamidino-2-phenylindole (DAPI; 0.02 mg/mL, ThermoFisher Scientific, USA), mounted and cover-slipped before being imaged with the Zeiss Axio Imager.Z1 Microscope. Percentage proliferative (Ki67 expressing cells) was calculated using equation 7.

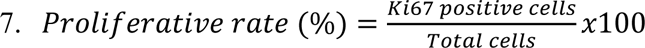

Growth patterns of MDA-MB-231 cells, the junction density per area was quantified using Angiotool software by measuring the number of cellular junctions in the explant area after staining cryosections with DAPI and Rhodamine phalloidin. Comparatively, the size of cellular aggregates and the percentage of the hydrogel occupied by cells (cell coverage, %) were assessed for MCF7 cells using ImageJ. Difference in analyses were due to the difference in morphology between these two cell types.

Hydrogels were divided into two regions based on where the cells were located: the “centre” and “edge” regions. The edge region was defined as the area within 200 µm of the outer top and sides of the hydrogels, while the centre region was defined as the remaining area that was 200 µm into the hydrogel and deeper (see Figure 2 for a visual representation of this division).

**Figure 2:**
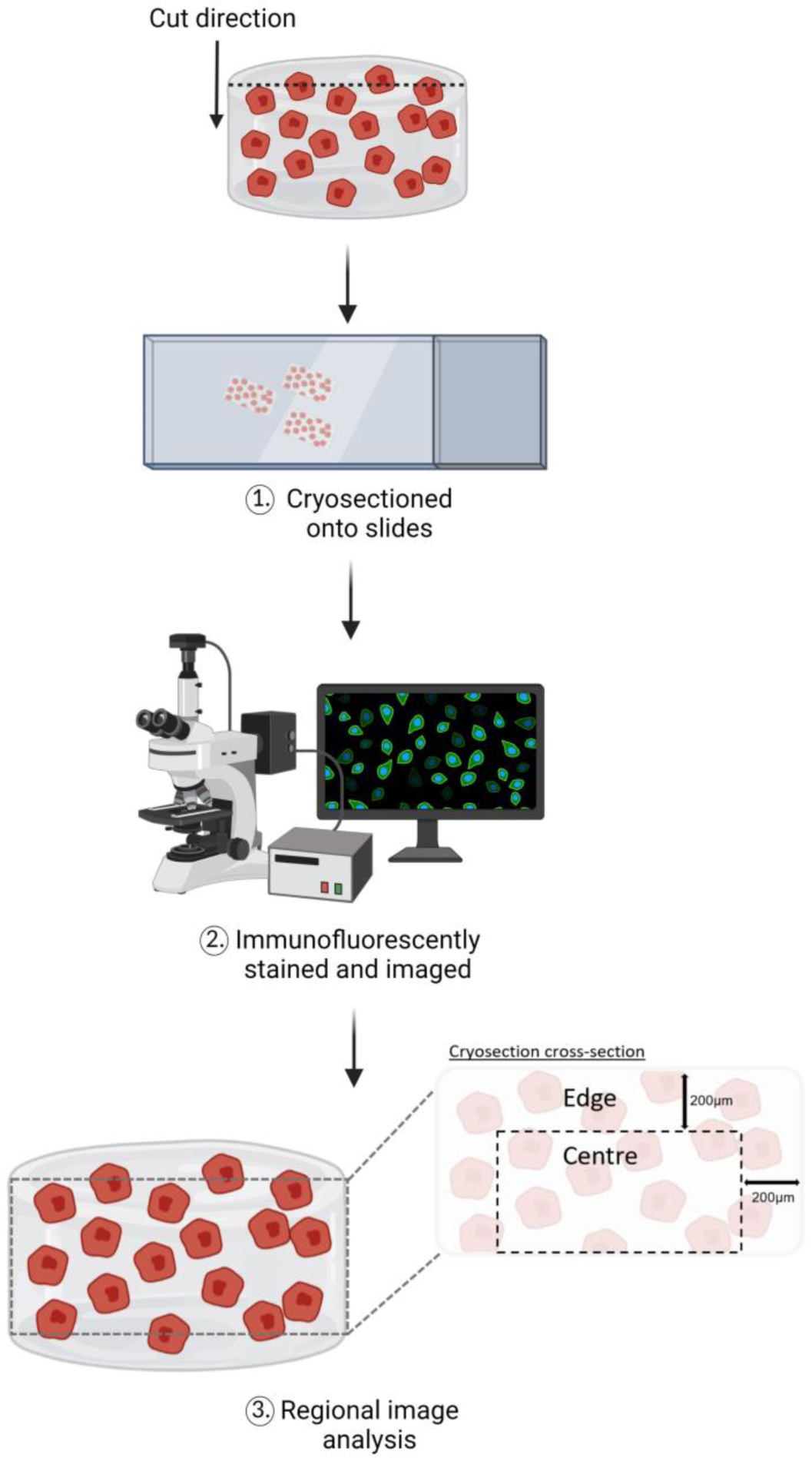
Schematic of regional image analysis in GelMA hydrogels. (1) Formalin-fixed and OCT embedded hydrogels were cryosectioned to produce 30 µm thick sections which were cut in a top-down manner onto adhesive microscope slides, (2) these sections were then immunofluorescently stained for proteins of interest and imaged on a fluorescence microscope. (3) Images were quantified based on the region of the hydrogel, which were separated into centre and edge regions. The edge region was defined as 200 µm from the outer top and side edges of the hydrogels and the centre region as the remaining region from 200 µm into the hydrogel and deeper. Image created with Biorender.

The ImageJ software analyser tool was used to quantify the relative expression of immunofluorescently stained proteins. The mean staining intensity (pixel density) was normalised to the number of cells within the field measured.

### Statistical analysis

Results were expressed as means ± standard deviation of the mean. GraphPad Prism software (version 9.2.0) was used to perform statistical analysis. Data resulting from a minimum of three independent biological repeats were evaluated, and statistical significance was determined using two-way ANOVA. A p-value of less than 0.05 was considered significant, and the degree of significance was denoted as follows: ns (not significant), * (p≤0.05), ** (p≤0.01), *** (p≤0.001), and **** (p≤0.0001).

## Results and Discussion

### Hydrogel characteristics and the impact of incorporating breast cancer cells

In previous studies, acellular hydrogels have been used to measure hydrogel properties^45–48^. However, it is well-established that encapsulation of cells can influence the photo-crosslinking process and introduces and extra component that could potentially affect the mechanical properties of the hydrogel through incomplete crosslinking.

Therefore, in order to create breast cancer cell-laden GelMA hydrogels of varying mechanical stiffness, 5 million cells/mL were encapsulated in hydrogels that contained 5%, 7.5%, and 10% GelMA macromer by weight. The concentration of 5 million cells/mL was selected as it falls within the reported range of concentrations used in 3D models, which typically range from 1 to 15 million cells/mL^22, 35, 47, 49–55^. The range of GelMA macromer concentrations (5%, 7.5%, and 10%) was selected based on previous studies that used these concentrations to produce hydrogels with mechanical stiffness similar to that of normal and malignant *ex vivo* patient breast tissues^13, 15^. The physical properties of the hydrogel were determined by measuring the hydrogel mass swelling ratio (q), soluble fraction (%), and compressive modulus (kPa).

The mass swelling ratio of hydrogels was measured to determine their swelling capacity. Cell-free GelMA hydrogel controls showed that increasing macromer concentration led to a decrease in mass swelling ratio (Figure 3a). The addition of MDA-MB-231 cells slightly increased the mass swelling of 5 wt% hydrogels, but not MCF7 cells. The mass swelling ratio of hydrogels was not significantly impacted by the addition of either cell line in 7.5 wt% and 10 wt% GelMA hydrogels. MDA-MB-231 and MCF7-laden hydrogels exhibited a similar decrease in mass swelling ratio as cell-free hydrogels with increasing macromer concentration.

**Figure 3:**
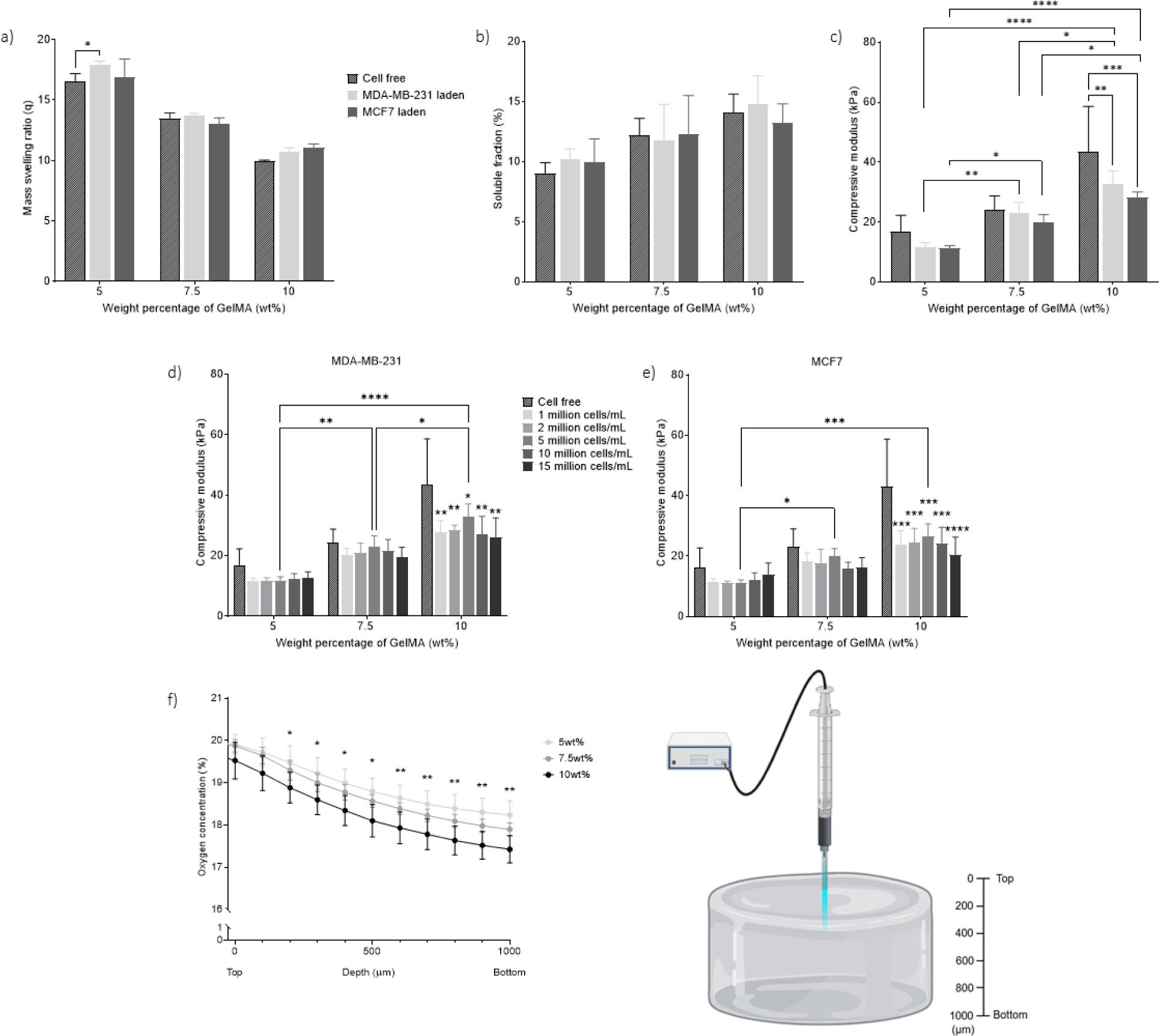
Impact of breast cancer cells on GelMA hydrogel properties: a) mass swelling ratio (q) (n=3), b) soluble fraction (%) (n=3) and c) compressive modulus (kPa) of cell-free and breast cancer cell-laden (5 million cells/mL) GelMA hydrogels (n=6) (hydrogels were 2 mm high). Breast cancer cells were encapsulated at 0, 1, 2, 5, 10 or 15 million cells/mL within GelMA hydrogels of 5, 7.5 or 10 wt%, d) MDA-MB-231 laden, e) MCF7 laden (n=3 for 1, 2, 10 and 15 million cells/mL data and n=6 for cell-free and 5 million cells/mL data). f) Measurement of oxygen concentration (%) throughout the depth of cell-free GelMA hydrogels of different stiffness (hydrogels were 1 mm high). Bars represent means ± SD, ns= not significant, *p≤0.05, **p≤0.01, ***=p≤0.001, ****p≤0.0001 by 2-way ANOVA with Tukey’s multiple comparisons test.

The soluble fraction of a hydrogel refers to the percentage of macromer that remains un-crosslinked and subsequently washes out of the hydrogel network. It was observed that the soluble fraction increased as the macromer concentration increased in cell-free GelMA hydrogels (Figure 3b). In comparison, the addition of breast cancer cell lines did not significantly impact the soluble fraction of GelMA hydrogels.

As expected, the compressive modulus of GelMA hydrogels, both cell-free and cell-laden, increased significantly with increasing macromer concentration (Figure 3c). The stiffness of 5 wt% and 7.5 wt% hydrogels did not change significantly with the addition of either MDA-MB-231 or MCF7 breast cancer cells. However, adding either cell type at a concentration of 5 million cells/mL reduced the stiffness of 10 wt% hydrogels significantly. There was no significant difference in the stiffness of 10 wt% hydrogels between those loaded with MDA-MB-231 or MCF7 cells.

In general, adding breast cancer cells at a concentration of 5 million cells/mL had little effect on the swelling and soluble fraction properties of GelMA hydrogels with 5, 7.5, and 10 wt% macromer, which is consistent with previous published literature^47, 56–58^. Only the 10 wt% hydrogels showed a decrease in stiffness with the addition of breast cancer cells at 5 million cells/mL.

To investigate whether the decrease in stiffness observed in 10 wt% GelMA hydrogels with 5 million cells/mL breast cancer cell encapsulation was influenced by cell concentration, a variety of breast cancer cell concentrations (0, 1, 2, 5, 10, and 15 million cells/mL) were introduced into GelMA hydrogels, and the corresponding compressive moduli evaluated (Figure 3d and e). The presence of MDA-MB-231 or MCF7 cells at various concentrations did not significantly affect the stiffness of 5 wt% and 7.5 wt% GelMA hydrogels. However, in the case of 10 wt% GelMA hydrogels, the addition of breast cancer cells at any concentration led to a significant reduction in stiffness (Figure 3d and e). Increasing cell concentration has been reported to reduce hydrogel stiffness^59, 60^, but the effect was not concentration-dependent in these GelMA hydrogels. In 10 wt% GelMA hydrogels, even low cell concentrations (1 million cells/mL) significantly reduced the stiffness. Interestingly, in 10 wt% hydrogels, stiffness initially increased with cell concentration (1 to 5 million cells/mL) but decreased at higher concentrations (10-15 million cells/mL). This suggests that increasing cell numbers initially contribute to hydrogel stiffness, but higher concentrations may impede crosslinking, leading to reduced stiffness. Of the investigated concentrations, the 5 million cells/mL condition in 10wt% GelMA hydrogels showed the least difference in stiffness to cell free controls. Based on these results, the 5 million cells/mL concentration was chosen for subsequent experiments.

At each GelMA macromer concentration, MDA-MB-231 laden hydrogels had a mean stiffness of 11.7 kPa (5 wt%), 22.9 kPa (7.5 wt%), and 32.7 kPa (10 wt%), while MCF7 laden hydrogels had a mean stiffness of 11.3 kPa (5 wt%), 19.9 kPa (7.5 wt%), and 28.2 kPa (10 wt%). Therefore, the range of GelMA hydrogel stiffnesses produced in this current study provided a panel of hydrogels which mimic the stiffness observed in breast tumour microenvironments with increasing grade (ranging from 10.4 in low grade, to 42.5 kPa in high grade IDC)^12, 13, 17, 18^, allowing assessment of the way different breast cancer cell types respond to these microenvironments.

Solid tumours are known to contain oxygen gradients, which can have significant effects on breast cancer phenotype and behavior, such as migration and invasion^6, 61–63^. To determine whether such gradients existed within the GelMA hydrogels, oxygen concentrations were measured in 5, 7.5, and 10 wt% hydrogels. Measurements were taken in a top-down manner throughout the depth of cell-free hydrogels, with 0 µm indicating the top and 1000 µm indicating the bottom of the hydrogel. Hydrogels with a height of 1 mm were assessed for direct comparison with subsequent cell response experiments. As expected, oxygen concentration decreased steadily with increasing hydrogel depth in all samples, with 10 wt% GelMA hydrogels exhibiting a significantly lower oxygen concentration than 5 wt% GelMA hydrogels at 200 µm and deeper (Figure 3f). The lowest concentration detected was 17% O₂ at the base of 10 wt% hydrogels, which does not reach levels that would be considered hypoxic in tumour tissue (<1% O₂)^7, 61, 64^. In the current study, a regional separation into the ‘centre’ and ‘edge’ regions of the hydrogel was employed to assess cellular response to the different O₂ concentrations that exist within these hydrogel regions (Figure 2).

### Breast cancer cellular response to culture in GelMA hydrogels of different stiffness

The response of both breast cancer cell lines to 5 wt% (soft), 7.5 wt% (medium) and 10 wt% (stiff) GelMA hydrogels was examined at day 1, 7, 14, and 21 of culture^36, 65–67^. MDA-MB-231 cells showed reduced viability in 7.5 wt% and 10 wt% hydrogels from day 7 onwards, while MCF7 cells only showed reduced viability in 10 wt% hydrogels at day 21 of culture (Figure 4a and b). Breast cancer cellular metabolic activity increased between day 1 and day 21 in all 5, 7.5 and 10wt% GelMA hydrogels (Figure 4c and d). Cell proliferation, as measured by DNA content, was greatest in both breast cancer cell lines when cultured in 5 wt% (soft) hydrogels. Previously reported studies which used 3D hydrogel cultures with a stiffness of around 5-15 kPa, yielded similar findings to the current study^47, 68, 69^. Specifically, these studies demonstrated that softer hydrogel conditions are conducive to cell functionality, as determined by assays measuring viability, metabolism, proliferation, or migration^47, 68, 69^. The 7.5 wt% (medium) stiffness hydrogels did not exhibit significant differences to soft or stiff hydrogels and were therefore not further examined in this study.

**Figure 4:**
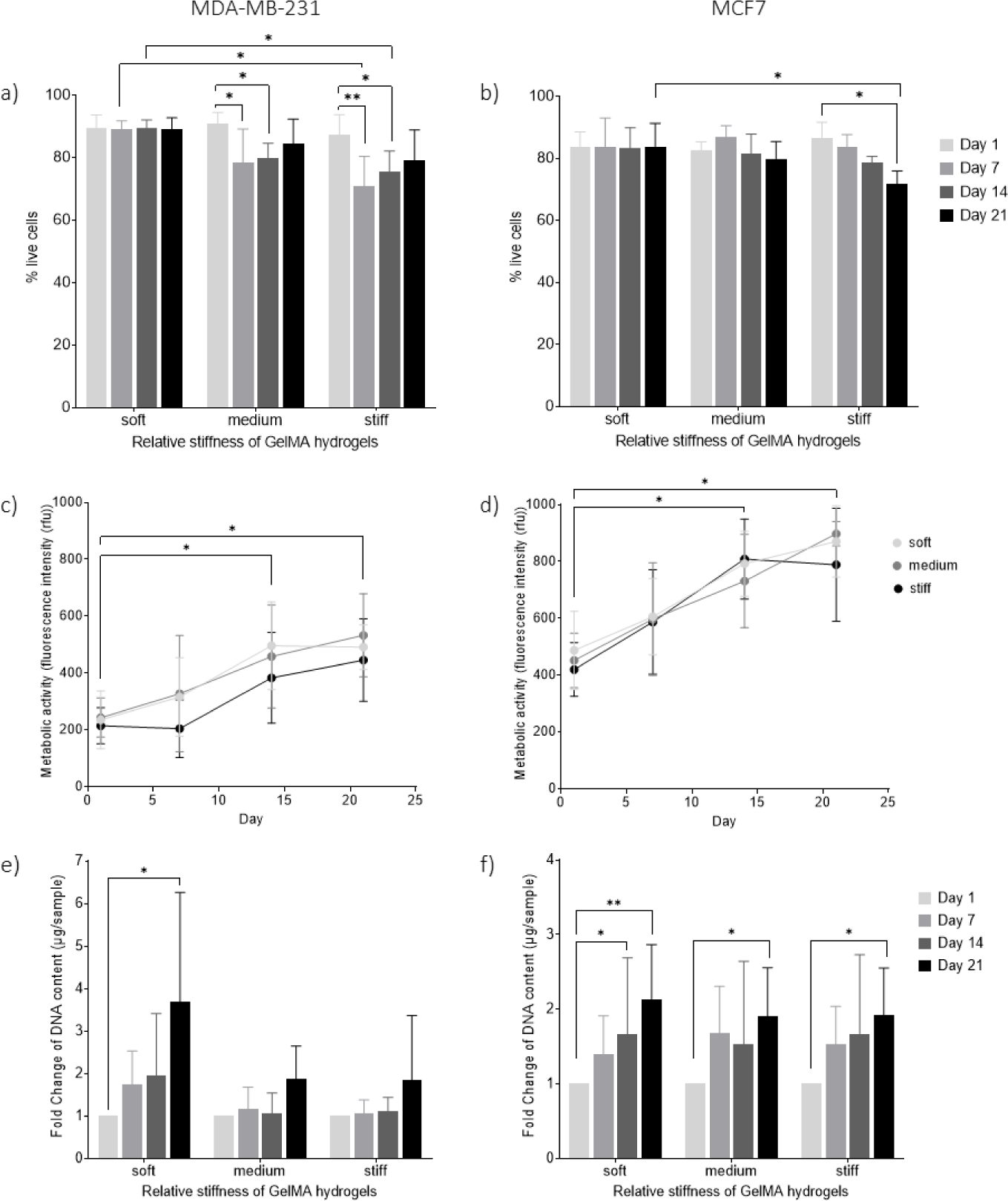
Breast cancer cell behaviour in GelMA hydrogels of different stiffness. MDA-MB-231 cells; a) cell viability (%) (n=5), c) metabolic activity (fluorescence intensity, rfu) (n=5) and e) fold change of DNA content (µg/sample) (n=3). MCF7 cells; b) cell viability (%) (n=3), d) metabolic activity (fluorescence intensity, rfu) (n=3) and f) fold change of DNA content (µg/sample) (n=3) in soft (5 wt%), medium (7.5 wt%) and stiff (10 wt%) GelMA hydrogels over 21 days. Hydrogels were 1 mm high. Bars or data points represent means ± SD, *=p≤0.05, **=p≤0.01, by 2-way ANOVA with Tukey’s multiple comparisons test. Rfu= relative fluorescence units.

MDA-MB-231 cells demonstrated a spindle-like, elongated morphology and formed connected networks within the GelMA hydrogels (Figure 5a). Angiotool software was used to quantify the junction density per area of MDA-MB-231 cells measured within 5 wt% (soft) and 10 wt% (stiff) hydrogels, separated into ‘edge’ and ‘core’ regions (Figure 5b). MDA-MB-231 cells developed the highest number of connections between cells over time within the edge region of hydrogels irrespective of hydrogel stiffness. Additionally, the centre region of soft hydrogels displayed greater capacity for formation of cellular junctions than the centre region of stiff hydrogels. This difference in MDA-MB-231 cell growth pattern suggests that these cells may be more proliferative within the edge region of these GelMA hydrogels.

**Figure 5:**
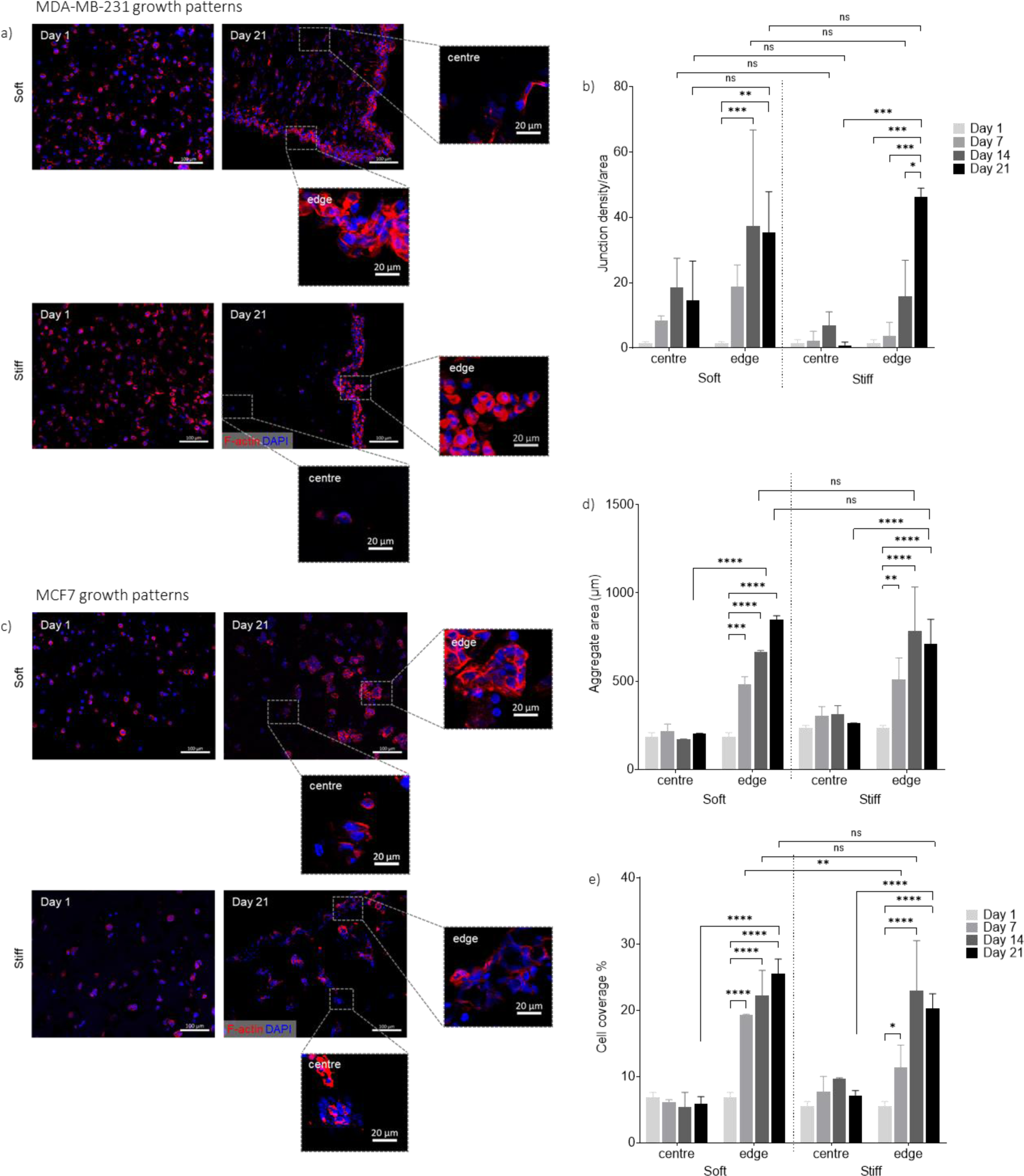
Breast cancer cell growth patterns within GelMA hydrogels of different stiffness. a) Representative images of MDA-MB-231 cells in soft (5 wt%) and stiff (10 wt%) GelMA hydrogels at days 1 and 21; zoomed inset images are of centre and edge regions of the hydrogel. b) Quantification of mean MDA-MB-231 cell junction density normalised to area (junction density/area) within the centre and edge regions of soft (5 wt%) and stiff (10 wt%) GelMA hydrogels at day 1, 7, 14 and 21 in culture. c) Representative images of MCF7 cells in soft (5 wt%) and stiff (10 wt%) GelMA hydrogels at days 1 and 21; zoomed inset images are of centre and edge regions of the hydrogel. d) Quantification of mean MCF7 aggregate area (µm) and e) MCF7 cell coverage (%), within the centre and edge regions of soft (5 wt%) and stiff (10 wt%) GelMA hydrogels at day 1, 7, 14 and 21 in culture. Hydrogels were 1 mm high. F-actin filaments were stained with rhodamine phalloidin (red) and nuclei with DAPI (blue). Bars represent means ± SD (n=3), ns=not significant, *=p≤0.05, **=p≤0.01, ***=p≤0.001, ****=p≤0.0001 by 2-way ANOVA with Tukey’s multiple comparisons test. Scale bars= 100 µm (20 µm on inset zoomed images).

The MCF7 cell line showed a clustering, aggregated growth pattern with cell-to-cell contact within the GelMA hydrogels (Figure 5c). The aggregate area (µm) and cell coverage percentage (%) were quantified within the centre and edge regions of soft and stiff hydrogels using ImageJ software (Figure 5d and e). These results suggest that the MCF7 cells in the edge region of hydrogels grow in a similar pattern (aggregate size and cell coverage %) regardless of hydrogel stiffness (Figure 5d and e). In addition, the centre region of both soft and stiff hydrogels showed similarly low aggregate size and cell coverage %, displaying no difference in response to stiffness. This highlights the ability of MCF7 cells to develop similar growth patterns despite hydrogel stiffness, but preferentially in the edge region of hydrogels, and this may be due to a greater proliferation within the edge region.

Ki67 is a protein expressed by cells at every stage of proliferation in the cell cycle and is absent in G0 or quiescent cells^70–72^. Based on the AJCC (American Joint Committee on Cancer) 2018 and the IKWG ((International Ki67 in Breast Cancer Working Group) 2021, the expression of Ki67 can be broadly separated into low (<10%), intermediate (10-20%), high (>20%) and very high (>30%) levels^73, 74^. Higher rates of proliferation are associated with poorer outcomes and Triple Negative breast cancers have been shown to exhibit much higher levels of Ki67 expression compared to Luminal A breast cancers^72, 75, 76^. The proliferative rate of MDA-MB-231 and MCF7 breast cancer cells were found to be similar when cultured in GelMA hydrogels, with both breast cancer cell lines displaying medium (10-20% Ki67 positive cells) to high (>20% Ki67 positive cells) proliferative rates. Results also showed that the proliferative rate of these cells was not greatly modified by the GelMA hydrogel stiffness, or region within the hydrogel (Figure 6).

**Figure 6:**
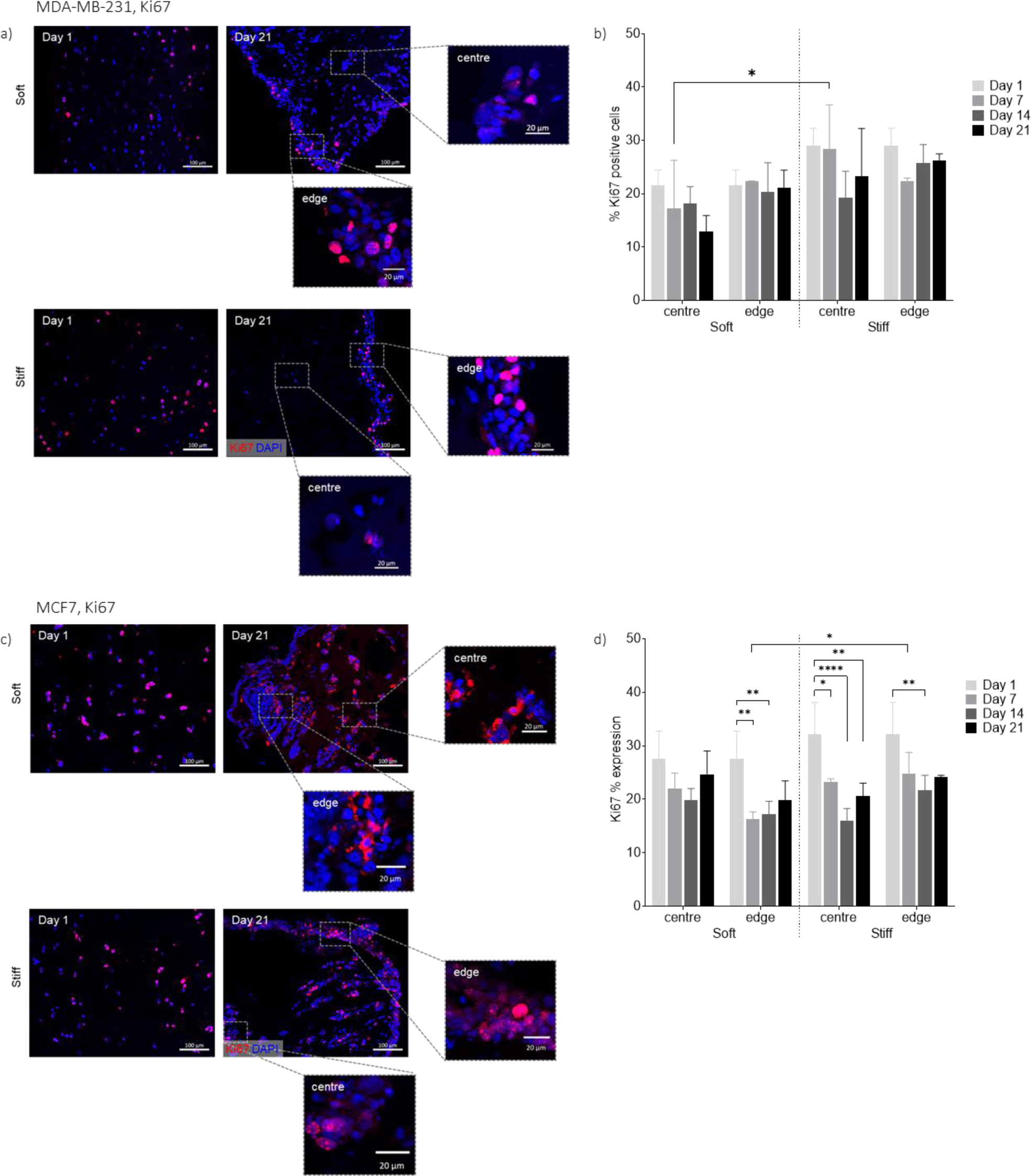
Breast cancer cell proliferative rate within GelMA hydrogels of different stiffness. a) Representative images of MDA-MB-231 cells in soft (5 wt%) and stiff (10 wt%) GelMA hydrogels at days 1 and 21; zoomed inset images are of centre and edge regions of the hydrogel. b) Quantification of the percentage (%) of proliferative (Ki67+) MDA-MB-231 cells within the centre and edge regions of soft and stiff GelMA hydrogels. c) Representative images of MCF7 cells in soft (5 wt%) and stiff (10 wt%) GelMA hydrogels at days 1 and 21; zoomed inset images are of centre and edge regions of the hydrogel. d) Quantification of the percentage (%) of proliferative (Ki67+) MCF7 cells within the centre and edge regions of soft and stiff GelMA hydrogels. Nuclei are stained with DAPI (blue) and proliferative cells stained for Ki67 (red). Bars represent means ± SD (n=3), *=p≤0.05, **=p≤0.01, ****=p≤0.0001 by 2-way ANOVA with Tukey’s multiple comparisons test. Scale bars= 100 µm (or, 20 µm on inset zoomed images).

The findings of this study suggest that the growth patterns observed in both MDA-MB-231 and MCF7 cells may not be solely due to cell proliferation, but could also be influenced by other mechanisms, such as cell migration within the hydrogel. MDA-MB-231 cells have been previously reported to be highly migratory in both 2D^77–79^ and 3D^53^ cultures, while MCF7 cells show limited migratory capacity in these systems^53, 80–82^. Additionally, the differences in oxygen concentration within the hydrogels, especially the lower oxygen concentration in the centre region, may also play a role in the observed growth patterns. Previous studies have shown that MDA-MB-231 cells migrate towards regions of higher oxygen concentration^83^, which may explain the increased junction density observed in the edge region of the hydrogel. However, this cannot fully account for the higher aggregate area and cell coverage percentage observed in the edge regions of the hydrogel in MCF7 cells, which are not known to be migratory. Therefore, further investigation is needed to elucidate the mechanisms by which different stiffness GelMA hydrogel microenvironments influence cellular growth patterns, including whether oxygen gradients in combination with increased hydrogel stiffness or other factors, such as nutrient gradients, are responsible for the observed differences.

### Impact of GelMA hydrogel stiffness on breast cancer cell EMT phenotype

Epithelial-mesenchymal transition (EMT) refers to a multifaceted process that enables cancer cells to enhance their migratory and invasive properties, ultimately leading to metastatic spread from the primary tumour^84–86^. This intricate process involves the downregulation of cell-to-cell adhesions and the upregulation of migratory and invasive phenotypes^87, 88^.

Results indicated that the mesenchymal MDA-MB-231 breast cancer cells did not exhibit changes in the expression of EMT-associated proteins E-cadherin (Figure 7a and b), N-cadherin (Figure 7e and f), or Vimentin (Figure 7i and j) after 21 days of culture in either soft or stiff GelMA hydrogels, which is not unexpected given they are characterised as having undergone partial-EMT^78, 89–91^. However, epithelial MCF7 breast cancer cells showed a decrease in the epithelial marker E-cadherin (Figure 7c and d), and increases in the expression of mesenchymal markers N-cadherin (Figure 7g and h) and Vimentin (Figure 7k and l) when cultured in stiff (10 wt%) GelMA hydrogels. This change in phenotype was only observed in MCF7 cells cultured in stiff (10 wt%) GelMA hydrogels, while no changes in E-cadherin, N-cadherin, or Vimentin expression were observed in MCF7 cells cultured in soft (5 wt%) GelMA hydrogels, suggesting increased stiffness of extracellular matrix in hormone receptor positive (Luminal A) breast cancers may promote breast cancer cell invasiveness via EMT^12–16^ (see Figure 8).

**Figure 7:**
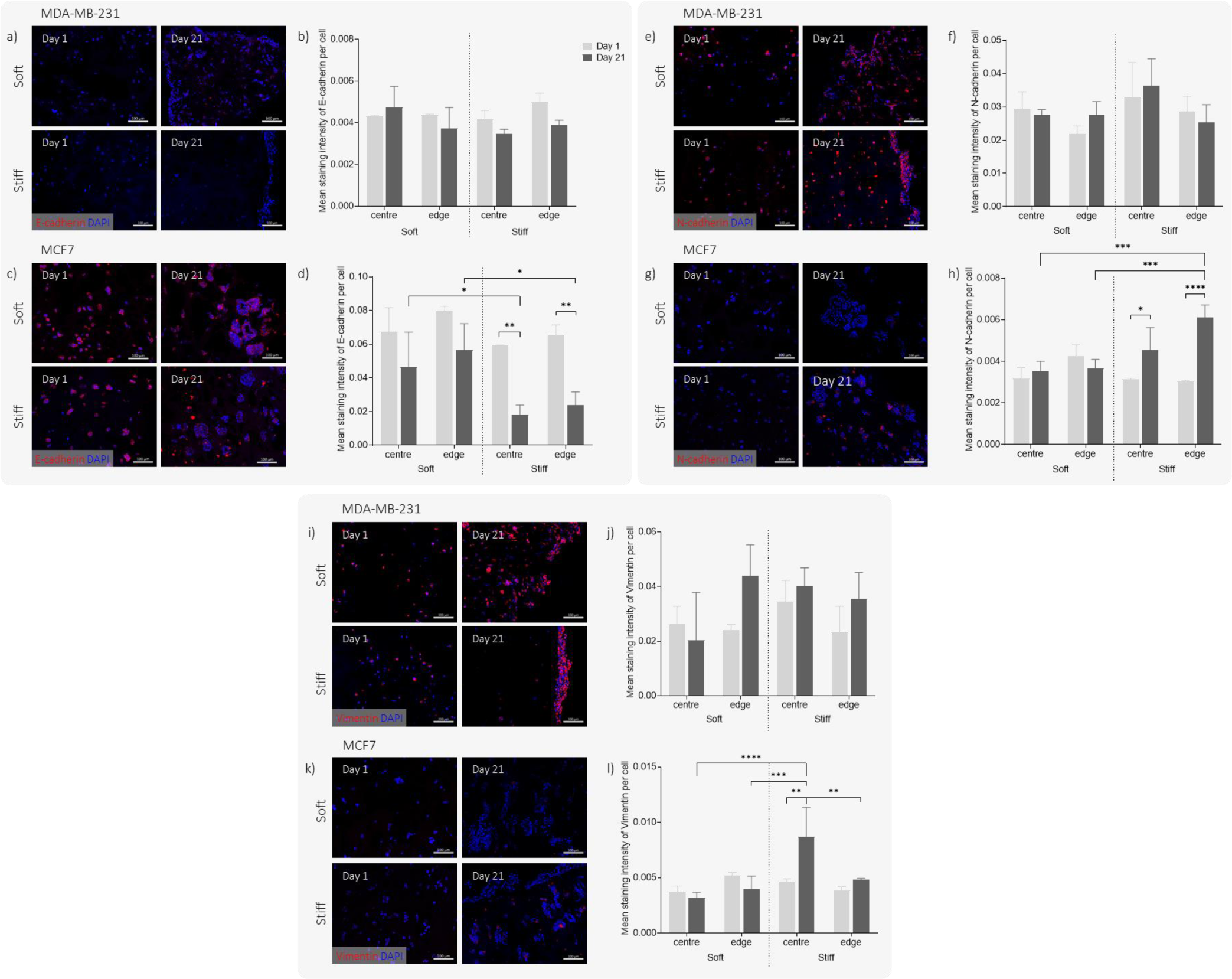
Breast cancer cell expression of E-cadherin, N-cadherin and Vimentin when cultured in GelMA hydrogels of different stiffness: a, e and i) Representative images of MDA-MB-231 cells stained for E-cadherin, N-cadherin and Vimentin, respectively, in soft (5 wt%) and stiff (10 wt%) hydrogels at days 1 and 21. b, f and j) Quantification of mean MDA-MB-231 E-cadherin, N-cadherin and Vimentin staining intensity, respectively, per cell in soft and stiff hydrogels at days 1 and 21. c, g and k) Representative images of MCF7 cells stained for E-cadherin, N-cadherin and Vimentin, respectively, in soft and stiff hydrogels at days 1 and 21. d, h and l) Quantification of mean MCF7 E-cadherin, N-cadherin and Vimentin staining intensity, respectively, per cell in soft and stiff hydrogels at days 1 and 21. Cells were stained with DAPI (nuclei, blue) and for E-cadherin, N-cadherin or Vimentin (red). Bars represent means ± SD (n=3), *=p≤0.05, **=p≤0.01, ***=p≤0.001, ****=p≤0.0001 by 2-way ANOVA with Tukey’s multiple comparisons test. Scale bars= 100 µm.

**Figure 8:**
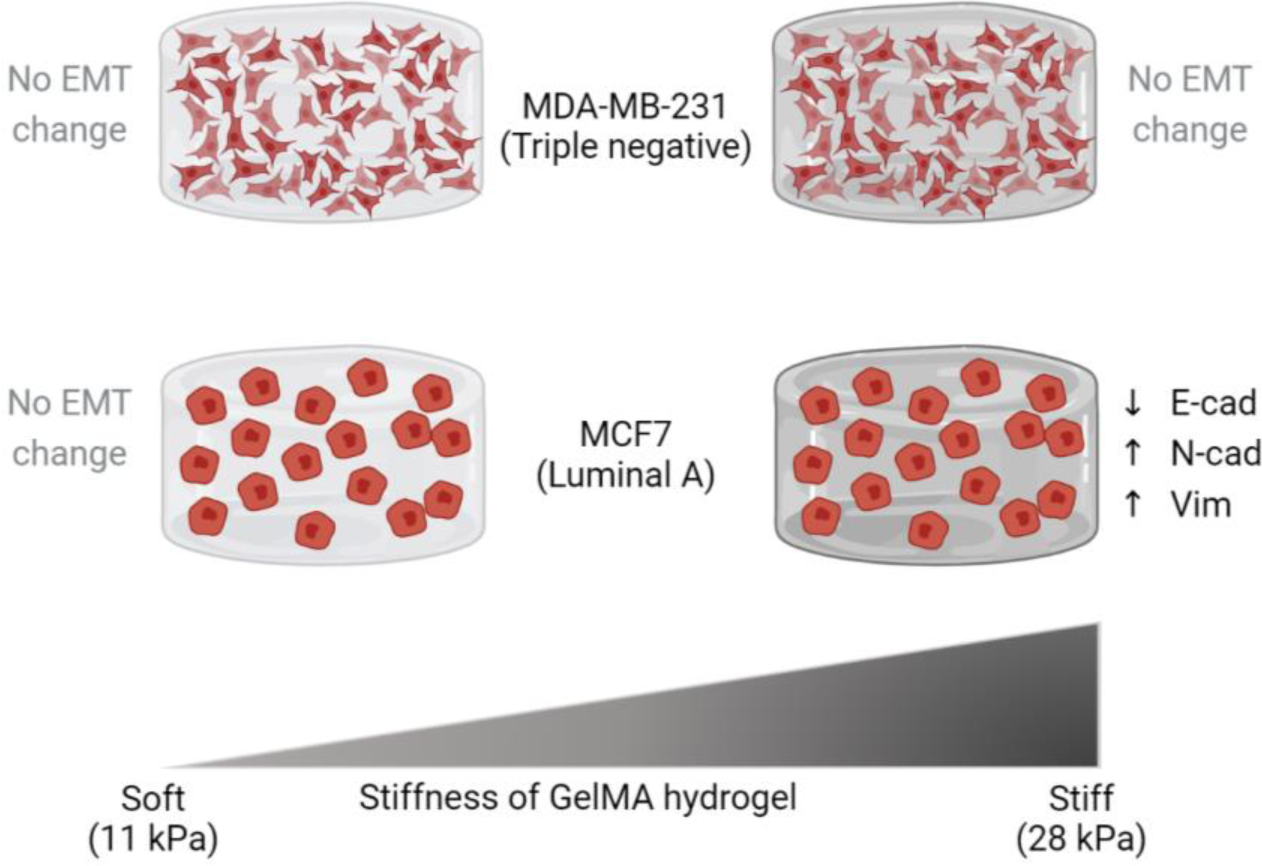
Graphical summary of major findings. Culture of MDA-MB-231 and MCF7 breast cancer cells within soft (11 kPa) GelMA hydrogels maintained the EMT phenotypes of both cell lines over a 21-day culture, while culture within stiff (28 kPa) GelMA hydrogels resulted in MCF7 cells undergoing EMT (downregulation of E-cadherin and upregulation of N-cadherin and Vimentin), revealing a breast cancer subtype specific response to matrix stiffness. Image created with Biorender.

When centre and edge regional data were separated, it was observed that the expression of E-cadherin or N-cadherin was not affected by the hydrogel region. However, the elevated expression of Vimentin at day 21 was observed only in the centre region of 10 wt% (stiff) hydrogels. Furthermore, the decrease in E-cadherin and increase in N-cadherin and Vimentin expression could facilitate the migratory behaviour of MCF7 cells, leading to their potential movement towards the edge region of stiff hydrogels. This could explain the larger aggregate area/cell coverage percentage detected in that region. However, it does not explain how the same trend in growth patterns was observed in soft hydrogels, where MCF7 cells maintain expression of E-cadherin and do not upregulate N-cadherin and Vimentin between day 1 and day 21. Previously published work using collagen gels showed that MCF7 cells underwent a reduction in E-cadherin expression and upregulation of EMT markers (Snail, Slug, TWIST1 and 2, Vimentin and β-catenin)^92^, however they performed no analyses of gel properties or mechanical stiffness. As a result, this is the first report to our knowledge of hydrogel stiffness differentially influencing the EMT status of encapsulated MCF7 breast cancer cells.

## Conclusions

This study explored the use of a 3D GelMA hydrogel model and tuneable visible light crosslinking system, in which the stiffness of the hydrogel can be adjusted by altering the macromer concentration. The study aimed to investigate the impact of extracellular matrix stiffness on breast cancer cell behaviour and phenotype over a 21-day culture period using GelMA hydrogels of different stiffness levels that are clinically relevant. The results showed that the response of breast cancer cells to varied hydrogel stiffness was dependent on the cancer subtype. MCF7 (Luminal A) cells exhibited an upregulation of mesenchymal markers N-cadherin and Vimentin, with downregulation of epithelial marker E-cadherin when cultured in stiff GelMA hydrogels (10 wt%; 28 kPa). However, stiff hydrogel culture (10 wt%; 33 kPa) had no impact on the expression of E-cadherin, N-cadherin, or Vimentin in MDA-MB-231 (Triple Negative) cells. Furthermore, maintaining the phenotype of both MCF7 and MDA-MB-231 breast cancer cells for 21 days was achievable by culturing them in soft hydrogels (5 wt%; 11.3-11.7 kPa). This study emphasizes the importance of carefully controlling hydrogel mechanical properties to preserve or modify breast cancer cell phenotype during 3D *in vitro* culture. Moreover, it provides a novel platform for studying the impact of extracellular matrix stiffness on the phenotype of various tumour-associated cell types, as well as the molecular mechanisms underlying cancer cell EMT, invasion and metastasis.

## Abbreviations

TME: Tumour Microenvironment
EMT: Epithelial-Mesenchymal Transition
IDC: Invasive Ductal Carcinoma
ECM: Extracellular Matrix
3D: three dimensional
2D: two dimensional
GelMA: Gelatin methacryloyl
UV: ultraviolet

## AUTHORS CONTRIBUTIONS

Conceptualisation: EP, KL, TW, MC; Data curation: JW; Formal analysis: JW; Funding acquisition: EP, KL, TW; Investigation: JW; Methodology: KL, EP, JW; Project Administration: EP, KL; Resources: KL, TW, EP; Supervision: KL, EP, MC, TW; Original draft preparation: JW; Review and editing: MC, EP, TW, KL. The first draft of the manuscript was written by JW and all authors commented on subsequent versions of the manuscript. All authors read and approved the final manuscript.

## ACKNOWLEDGEMENTS

This research was funded by the University of Otago (UORG), Maurice Wilkins Centre (MWC), and the Cancer Society NZ Canterbury West Coast Division.

## CONFLICTS OF INTEREST

The authors state no conflict of interest.

---

**Supplementary Figure:**
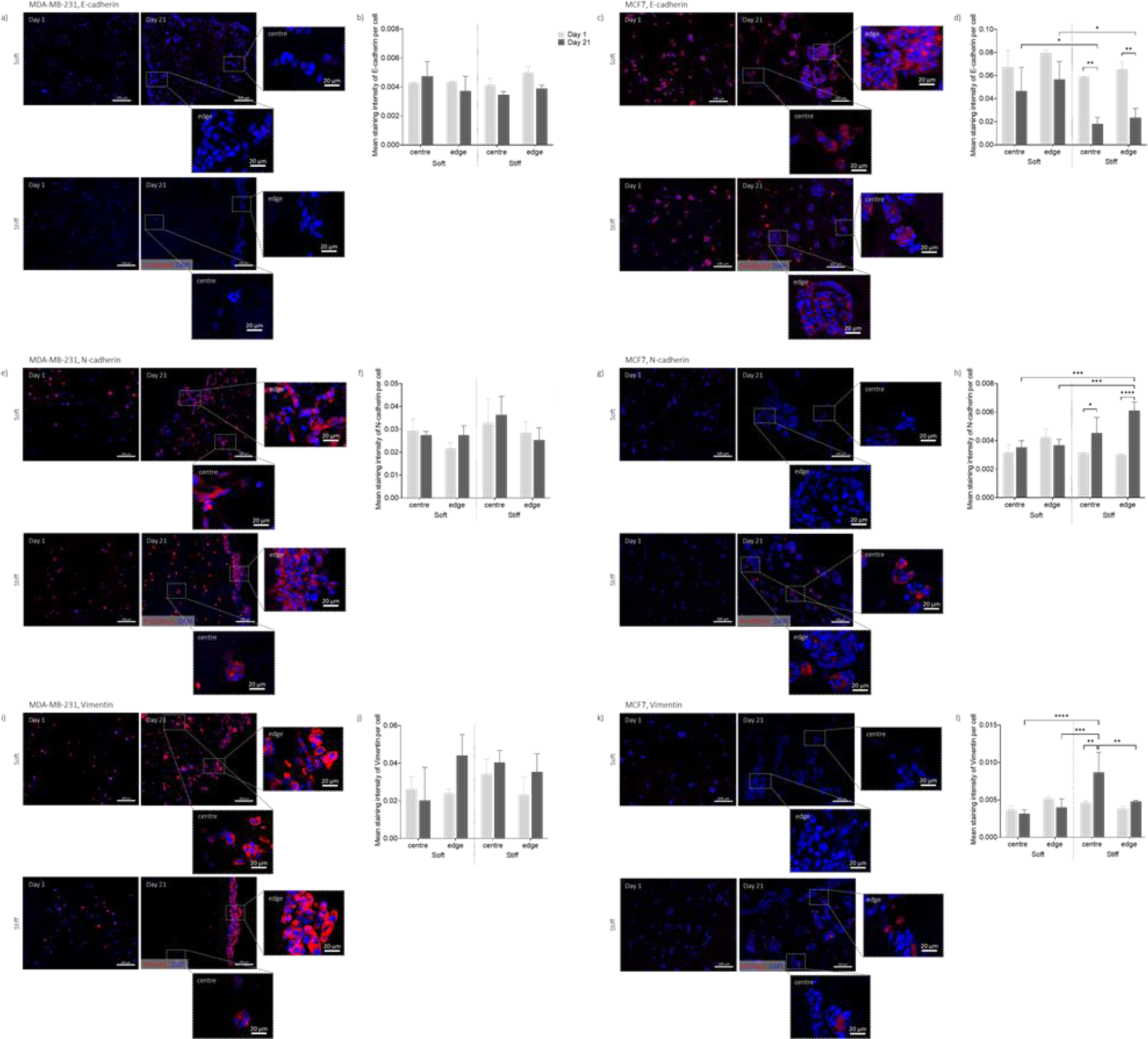
Additional to Figure 7, with regional zoomed inset images. Breast cancer cell expression of E-cadherin, N-cadherin and Vimentin when cultured in GelMA hydrogels of different stiffness: a, e and i) Representative images of MDA-MB-231 cells stained for E-cadherin, N-cadherin and Vimentin, respectively, in soft (5 wt%) and stiff (10 wt%) hydrogels at days 1 and 21; zoomed inset images are of centre and edge regions of the hydrogel. b, f and j) Quantification of mean MDA-MB-231 E-cadherin, N-cadherin and Vimentin staining intensity, respectively, per cell in soft and stiff hydrogels at days 1 and 21. c, g and k) Representative images of MCF7 cells stained for E-cadherin, N-cadherin and Vimentin, respectively, in soft and stiff hydrogels at days 1 and 21; zoomed inset images are of centre and edge regions of the hydrogel. d, h and l) Quantification of mean MCF7 E-cadherin, N-cadherin and Vimentin staining intensity, respectively, per cell in soft and stiff hydrogels at days 1 and 21. Cells were stained with DAPI (nuclei, blue) and for E-cadherin, N-cadherin or Vimentin (red). Bars represent means ± SD (n=3), *=p≤0.05, **=p≤0.01, ***=p≤0.001, ****=p≤0.0001 by 2-way ANOVA with Tukey’s multiple comparisons test. Scale bars= 100 µm (or, 20 µm on inset zoomed images).

## Notes

### Competing Interest Statement

The authors have declared no competing interest.

